# Task-Based Spinal fMRI using a Median Nerve Stimulation Reveals Functional Alterations in Degenerative Cervical Myelopathy

**DOI:** 10.1101/2025.09.17.675603

**Authors:** Grace Haynes, Fauziyya Muhammad, Kenneth A. Weber, Ali F. Khan, Sanaa Hameed, Hakeem J. Shakir, Michael Rohan, Lei Ding, Zachary A. Smith

**Affiliations:** Stephenson School of Biomedical Engineering, University of Oklahoma, Norman, OK; Department of Neurosurgery, University of Oklahoma Health Sciences Center, Oklahoma City, OK; Division of Pain Medicine, Stanford University School of Medicine, Palo Alto, CA; Laureate Institute for Brain Research, Tulsa, OK

**Keywords:** Spinal cord functional MRI, degenerative cervical myelopathy, biomarkers, median nerve stimulation

## Abstract

Functional spinal cord MRI (fMRI) is an emerging technique for evaluating sensorimotor responses in both healthy and disease states. However, degenerative cervical myelopathy (DCM), a non-traumatic, age-related pathology characterized by spinal cord compression, often leads to motor and sensory impairments that make standard task-based fMRI paradigms challenging to perform or execute. To overcome this, we implemented an fMRI paradigm using direct electrical stimulation to the median nerve to evoke neural responses in both motor and sub-motor spinal cord pathways. In this study, 23 DCM and 23 aged-matched healthy controls (HC) underwent two task-based spinal cord fMRI sessions with direct, median nerve stimulation to the right upper limb. Motor thresholds were determined on an individual basis by a thumb/finger twitching, and sub-motor thresholds were set to be 15% underneath motor with no visible twitching. Blood Oxygenation Level Depended (BOLD) signals were analyzed across the C5-C8 cervical spinal cord segments. HCs demonstrated increased activation during motor vs sub-motor stimulation (1155 vs 496 voxels; +7.1%), whereas DCM patients showed reduction in activation with higher stimulation (1028 vs 1220 voxels; -2.2%). This activation pattern was consistent across the cervical spinal segments, with HCs peaking at C7 (sub-motor: 328 voxels, motor: 608 voxels), which is consistent with median nerve input for forearm and hand. While DCM responses were altered, with sub-motor differentially maximized at C6 (406 voxels) and abnormal recruitment at C5 (112 voxels) and C8 (198 voxels). Spatially, HC responses were focused in the gray matter (GM) and bilateral, whereas DCM responses showed reduced GM localization and were more ipsilateral. Furthermore, we saw spikes in sub-motor DCM deactivation at the overall (3032 voxels) and C6 root level (1500 voxels) that were greater than the other conditions. This study demonstrates the feasibility and effectiveness of median nerve stimulation as a task-based paradigm for assessing spinal cord function in those with spinal cord compression. Group level differences between DCM patients and age-matched HCs showed spatial and quantitatively distinct patterns across stimulation thresholds and spinal segments. Our findings suggest disrupted sub-motor and motor activation in the DCM group, highlighting the use of this approach as a functional biomarker to capture disease specific alterations.

## 1 Introduction

Degenerative cervical myelopathy (DCM) is a common condition that is caused by progressive degenerative compression of the cervical spine. DCM has been known to impact descending white matter (WM) pathways (Cloney et al., 2018; Haynes et al., 2024; Hopkins et al., 2018), impairing strength, coordination, and balance (Mechas et al., 2025; Muhammad et al., 2023; Sharma et al., 2025). As the disease progresses, this injury can begin to spread the gray matter (GM) horns (Smith et al., 2020) and may result in sensorimotor impairment and irreversible neural tissue injury (i.e. myelomalacia) (Al-Mefty et al., 1988; Mehalic et al., 1990; Tsuchiya et al., 2003).

However, evaluating injury to the descending WM pathways and developing biomarkers capable of quantifying the ongoing WM damage has been the primary focus of most DCM imaging studies. Therefore, an investigation into the impact of significant chronic injury on the GM horns is needed to evaluate the contributions it has to the impairment of sensorimotor functions occurring with progressive injury.

One technique that could provide a way of evaluating the injury to the spinal cord and its resulting impact on functionality is functional MRI (fMRI), which can employ sensory (e.g. tactile or thermal stimulation) (Weber et al., 2020; Weber et al., 2018) and motor (e.g. fist clenching or finger abductions) (Kinany et al., 2023; Landelle et al., 2021) tasks to interrogate the functional pathways. These paradigms have been primarily used in subjects with healthy spinal cords, but they do also serve as an important foundation for applying task-based fMRI methods in cervical spinal diseases to study alterations in functional pathways (Haynes et al., 2023). However, only resting-state fMRI has been applied to study DCM induce injury, demonstrating that the Amplitude of Low Frequency Fluctuation (ALFF) in the cervical cord is associated with myelopathy severity (Liu et al., 2016). Clinical impairments associated with DCM manifestation (Hameed et al., 2024; Muhammad et al., 2023) means that the performance of task-based paradigms normally applied in healthy spinal cords can be negatively impacted.

Therefore, in this study, we apply direct electrical stimulation to the median nerve as a task-based fMRI paradigm to avoid the challenges that arise with subject-driven motor tasks and evaluate the functional pathways in the cervical spine of DCM patients by activating spinal cord reflex arcs. To date, median nerve stimulation has not been applied in DCM fMRI studies, but has been used in several fMRI papers involving healthy controls (Backes et al., 2001; Wilmink et al., 2003; Xie et al., 2007) where it was found to induce activation in the lower cervical spine (Backes et al., 2001; Wilmink et al., 2003; Xie et al., 2007) comparable to fist clenching (Backes et al., 2001; Wilmink et al., 2003). Direct stimulation is also a common electrodiagnostic tool used to study and diagnose functional changes in DCM patients (Dvorak et al., 2003) and median nerve stimulation specifically has been established to induce activation in areas commonly impacted by DCM compression (i.e. lower cervical spine) (Muhammad et al., 2024; Vallotton et al., 2021). Because this chronic compression can lead to progressive injury of the motor and sensory GM horns, we seek to demonstrate that median nerve stimulation can reveal functional impairment of sensorimotor pathways in DCM. Specifically, we focus on segmental recruitment in the lower cervical cord, recruitment differences between sub-motor and motor activation, and the spatial distribution of responses across GM, WM, and lateralized pathways.

## 2 Methods

### 2.1 Participants

DCM patients (n = 23, 17 females) were recruited from OU Health Adult Neurosurgery Clinic in Oklahoma City. All patients were evaluated independently by spine neurosurgeons (ZAS, HJS). Healthy control participants in this study (n = 23, 15 females) were simultaneously recruited from the Laureate Institute for Brain Research (LIBR) in Tulsa, OK and the University of Oklahoma Health Sciences Center (OUHSC). The HC (49.7 ± 2.23 years) and DCM (54.4 ± 2.03 years) groups were age-matched and had no significant difference between them (p = 0.12). All participants were right-handed. All participants were scanned between March 2021 and May 2024 and signed an informed consent document before data collection began. Details about this cohort, including inclusion and exclusion criteria, can be found in Muhammad, *et al*. (2025) (Muhammad et al., 2025). All imaging experiments performed in this study were approved by the University of OUHSC and LIBR institutional review boards (IRB #12068) and performed according to IRB regulation guidelines.

### 2.2 MRI Data Acquisition

Structural and functional cervical spine MRI data were collected with a Signa 3T MR750 (GE Healthcare, Milwaukee, WI) scanner equipped with a 32-channel NovaCSpine coil at LIBR. Participants laid supine in the scanner with no visual stimulation and were instructed to remain still while undergoing scanning. Isotropic T2-weighted sagittal and axial CUBE images (TR = 2500ms; TE = 120ms; FOV = 256 x 256mm; Thickness = 0.8mm; Resolution = 0.8 x 0.8mm) and a multi-echo gradient-echo T2*-weighted (TR = 600ms; FOV = 224 x 224mm; Thickness = 5mm; Resolution = 0.5 x 0.5mm) scan were collected according to the generic spine protocol (Cohen-Adad et al., 2021). Functional MRI (fMRI) scans were acquired with single-shot echo-planar imaging (SS-EPI) (TR = 2000ms; TE = 29.5ms; Flip Angle = 90 degrees; FOV = 128 x 128mm; Thickness = 5mm; Resolution = 1 x 1mm) covering the C3-T2 vertebral levels and the field of view was centered at C5.

### 2.3 fMRI Task Design: Stimulation Paradigm and MRI Setup Conditions

Ag/AgCl electrodes were placed on the center, inner wrist (cathode) and above the inner elbow (anode) of the subject’s right upper limb (Figure 1B). This placement of electrodes was based on the anatomy of the median nerve, which runs beneath the bicipital aponeurosis at the elbow and the carpal tunnel at the wrist (Mazurek & Shin, 2001). Due to individual neurological differences, stimulation thresholds could not be generalized for all subjects and were, therefore, determined individually. To do this, single monophasic electrical pulses were delivered at intervals of 0.5mA until a consistent involuntary twitching in the thumb or forefinger was observed. The amplitude of the pulse was then decreased by 0.1mA until the strength of the stimulation was only capable of inducing a minimally observable finger twitching (motor threshold). The sub-motor threshold was set at 15% below motor threshold to evoke sub-motor stimulation.

**Figure 1.**
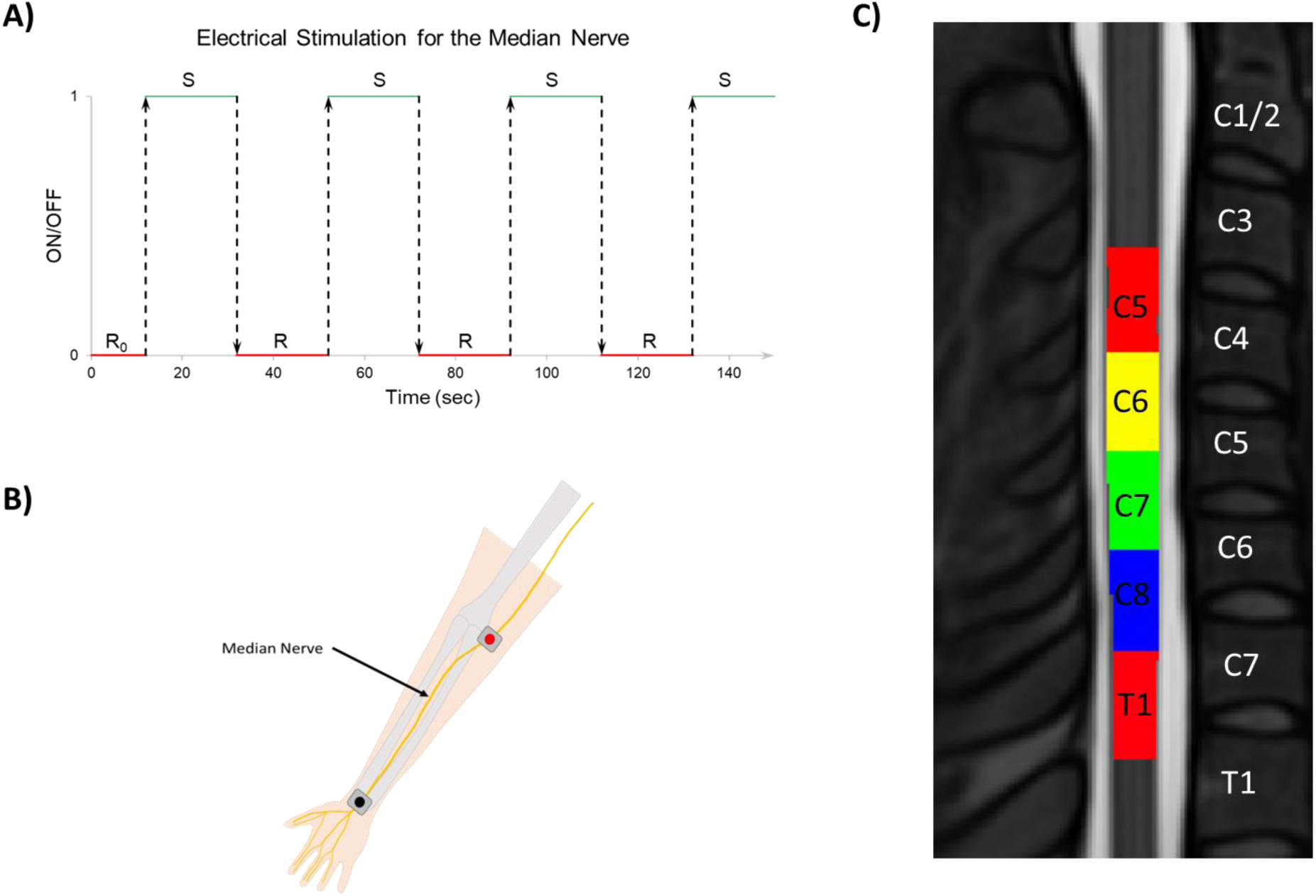
Overview of the spinal cord task-based fMRI stimulation paradigm and segmental localization. (A) Stimulation block design showing the timing of electrical median nerve stimulation. The paradigm begins with a 12-second initial rest period (R0), followed by alternating 20-second stimulation (S) and rest (R) defined time period across four cycles. (B) Schematic showing the surface localizations of median nerve in the forearm, illustrating the placement of surface electrodes. (C) Anatomical segmentation and masks of the cervical (C5-C8) and upper thoracic (T1) spinal segments using the PAM50 template. Colored masks correspond to spinal cord segments from C5 to T1, used for group level voxel extraction.

These thresholds were then applied in 2000μs pulses at 1 Hz for twenty seconds during the course of an 8 minutes, 12 seconds fMRI run, where they alternated with twenty second (i.e., rest) period until the end of the scan (Figure 1A). Each subject underwent two fMRI runs, which used the sub-motor and the pre-determined motor threshold stimuli respectively. To deliver consistent stimulation pulses during functional MRI scans, a series of trigger pulses were generated with a timed code synchronized with the beginning of the scan. Digital signals from a laptop were delivered to the input port of a high voltage, constant current stimulator (Digitimer DS7A), which outputted to a filtered 9-pinned series port (BIOPAC MRIRFIF DSUB-9) connected to an MRI compatible electrode cable (BIOPAC MECMRI-1) inside of the MRI suite. Anode and cathode leads then delivered the pulse to the electrodes located on the subject’s dominant (i.e. right) arm. Figure 2 demonstrates how the equipment was set up in an MRI environment.

**Figure 2.**
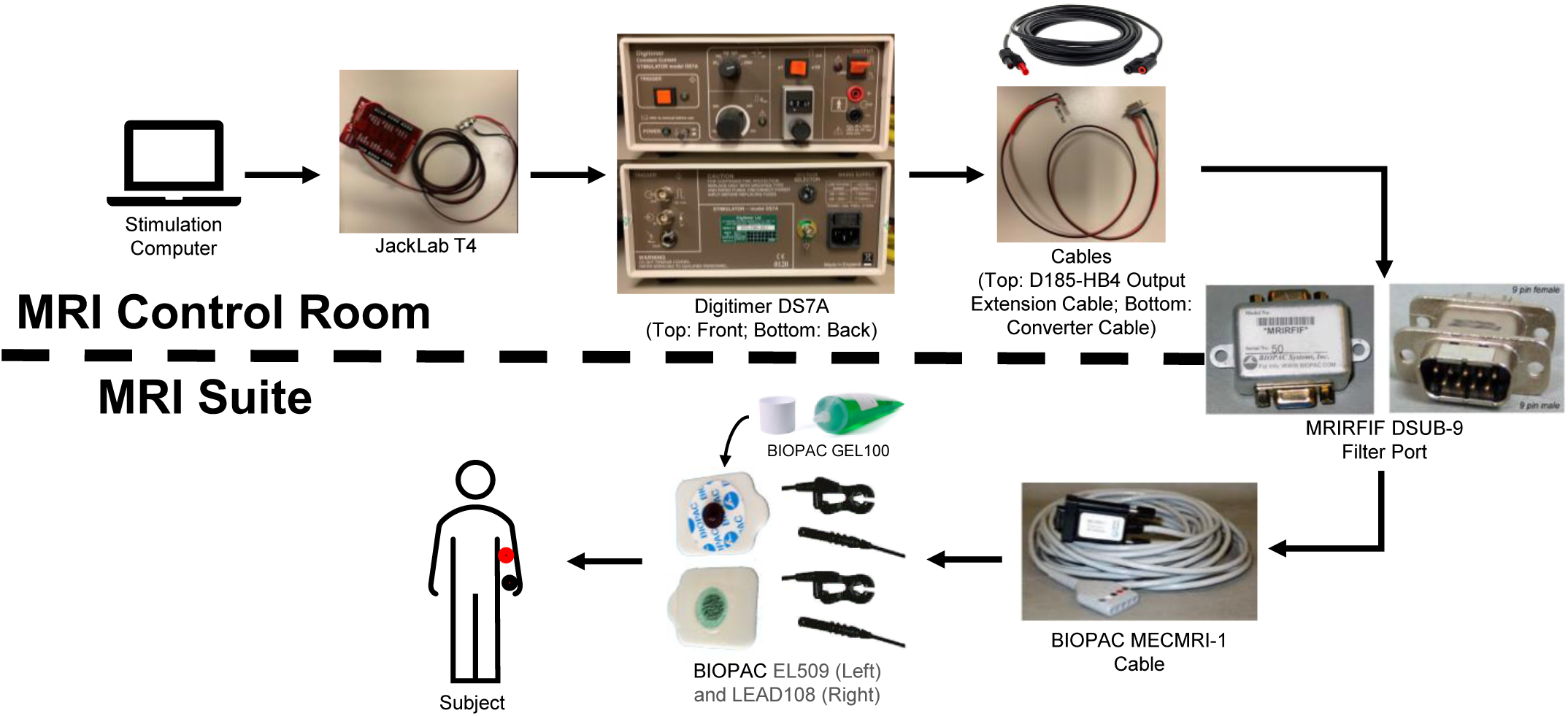
MRI-compatible electrical stimulation setup for median nerve stimulation. The stimulation protocol begins in the MRI control room, where the stimulation computer delivers the stimulation commands through JackLab T4 device. This device interfaces with a Digitimer (DS7A constant current stimulator (top: front view, bottom: rear view with connections). Stimulation pulses are routed through a series of output and converter cables that passed through an MRIRFIF DSUB-9 filter port into the MRI suite. Inside the MRI suite, filtered signals are transmitted via a BIOPAC MECMRI-1 cable to surface electrodes (BIOPAC EL509 and LEAD108) placed over the median nerve on the subject’s forearm. Conductive gel (BIOPAC GEL100) is applied to ensure correct contact and signal delivery during scanning.

### 2.4 MRI Processing

Spinal Cord Toolbox (SCT) (v.7) (De Leener et al., 2017) was used for structural image preprocessing for individual subjects. Spinal cord segmentation was performed using the *sct_deepseg_sc* function (Gros et al., 2019) and registered to the PAM50 template using *sct_register_to_template* (Ullmann et al., 2014) (Figure 1C). The structural spinal cord was registered to the PAM50 template using *sct_register_to_template* (De Leener et al., 2018). Lastly, the PAM50 templates were warped to the native space using *sct_warp_template*. Spinal cord segmentation was also created for the T2* images using *sct_deepseg_sc* (Gros et al., 2019). Gray matter (GM) segmentation was also extracted from the T2* images using *sct_deepseg_gm*, which was used in combination with the cord mask to create a white matter (WM) mask. The T2* image and white matter segmentation were then registered to the PAM50 template and warped to the native space using *sct_register_multimodal* and *sct_warp_template*. Further details about how anatomical images with compressed spinal cords were preprocessed can be found in Muhammad, F., et al. (2025) (Muhammad et al., 2025).

### 2.5 Functional Pre-Processing

The fMRI data was preprocessed using a combination of SCT (v.7) (De Leener et al., 2017) and FMRIB Software Library (FSL) (Jenkinson et al., 2012). The preprocessing steps follow similar description in previous studies using similar methodologies (Banerjee et al., 2025; Oliva et al., 2025; Weber et al., 2016a, 2016b). These preprocessing include:

#### 2.5.1 Motion Correction

Motion correction was performed in two stages using a custom 2D slice-wise registration pipeline optimized for spinal cord fMRI (Weber et al., 2020; Weber et al., 2016a, 2016b). First, a temporal mean image was generated from the raw BOLD time series using *sct_maths,* and the spinal cord was segmented using *sct_deepseg_sc.* A mask of the spinal canal was also created using *sct_propseg -CSF* (De Leener et al., 2015; Oliva et al., 2025), and the two masks were combined and dilated (disk shape, 5 voxels, 2D) to define a robust registration mask.

In the first motion correction step, each volume was registered slice-by-slice to the middle volume of the functional time series to limit motion correction the region within the registration mask. In the second step, the temporally averaged motion-corrected image from the first pass was used as the new reference volume, and all volumes were realigned to it using the updated registration mask. This two-pass approach improves motion correction by refining alignment across both time and anatomical structure, accounting for non-rigid, slice-specific motion frequently observed in spinal cord data (Weber et al., 2016a, 2016b).

#### 2.5.2 Segmentation and Registration to Template Space

Following motion correction, spinal cord segmentation was performed on the motion-corrected mean functional image. A subject-specific spinal cord mask was obtained by applying SCT’s EPI-trained deep learning segmentation tool (*sct_deepseg sc_epi*), which is optimized for EPI images (Banerjee et al., 2025). The spinal canal was then segmented using *sct_propseg -CSF* (De Leener et al., 2015), and this mask was dilated in-plane (disk, 3 voxels) to define the CSF region surrounding the cord.

For spatial normalization, functional data were registered to the PAM50 T2* spinal cord template using *sct_register_multimodal.* The registration steps included (1) slicewise segmentation-based alignment to the center of mass (i.e. *centermass*), (2) non-linear symmetric normalization alignment using b-splines (i.e. *bsplinesyn*), and (3) intensity-based refinement using non-linear symmetric normalization (i.e *syn*) with cross-correlation. Subject-specific warps derived from high-resolution T2* structural scans were used to initialize the registration. Finally, *sct_warp_template* was used to warp PAM50 template labels into functional space, enabling spinal level localization and group-level comparisons. For further details, please see the GitHub link in the “Data and Code Availability” section.

#### 2.5.3 Physiological Noise Modeling

Physiological denoising was performed using the RETROICOR-based approach, which modeled and removed cardiac and respiratory artifacts from the fMRI data using the phase regressors derived from the physiological recordings (Glover et al., 2000).

We also extracted the time courses from WM and CSF masks. The PAM50 T2* template was registered to each subject’s motion-corrected mean functional image using the *sct_register_multimodal* and *sct_warp_template*. The PAM50 WM mask and the segmented CSF canal (*sct_propseg -CSF* (De Leener et al., 2015)) were warped and binarized to extract mean time series from each axial slice. These were concatenated into regressors. These physiological regressors were entered into the denoising model implemented using the FSL’s FMRI Expert Analysis Tool (FEAT) (Weber et al., 2016a, 2016b).

#### 2.5.4 Slice-timing Correction and Smoothing

Denoised data were corrected for slice-timing offsets using FSL’s *slicetimer* tool, and the resulting volumes were warped to PAM50 template space using *sct_apply_tranfo* and previously estimated warp fields. Voxels falling outside of the automated spinal cord segmentation mask space were then removed using a spinal cord mask warped to template space. The images were then smoothed using FSL’s *susan* with a 2mm Full Width at Half Maximum (FWHM) Gaussian kernel (Smith & Brady, 1997). A representative example of signal stability in the spinal cord (via temporal signal to noise ratio) can be seen in Supplemental Figure 1.

### 2.6 Statistical Analysis

#### 2.6.1 First-Level Analysis

Subject-level analysis was performed in FSL’s FEAT (Woolrich et al., 2001) using a General Linear model (GLM) framework based on a square-shaped ON/OFF stimulation paradigm (Figure 1A) convolved with a gamma hemodynamic response function. A high-pass filter (cut-off: 0.01Hz) was then applied to remove low-frequency noise. DVARS-derived motion outliers (i.e. volumes where the difference between root mean squared values are greater than the 100^th^ percentile) were used as covariates of no interest. Temporal autocorrelation was modeled and corrected using FMRIB’s Improved Linear Model (FILM) (Woolrich et al., 2001). Voxel-wise activation was assessed using z-statistics and a threshold of z > 2.3 (uncorrected p-value of < 0.05) was applied to define activation.

#### 2.6.2 Group Level Analysis

Group-level statistical analysis was performed using the Higher-Level Analysis functionality of FSL’s FEAT tool (Woolrich et al., 2004). Analyses were performed for each contrast using GLMs implemented in FSL, with variance modeled at the group level. Three statistical tests were applied to compare group level activation results. The first tests were one-sample t-tests to detect whether BOLD signal changes were significantly different from zero within each group (HC and DCM) and condition (sub-motor and motor stimulations). These tests identify voxels that exhibited positive (BOLD z-statistic > 0) or negative (BOLD z-statistic < 0) activation. The second tests were paired two-sample t-tests applied within each group (HC and DCM) to compare motor vs sub-motor stimulation, identifying condition-driven differences in BOLD activation (i.e., motor > sub-motor and sub-motor > motor). The final approach applied was unpaired two-sample t-tests to compare activation between HCs and DCM groups during both stimulation conditions (i.e. HC > DCM and DCM < HC), assessing for disease-related differences. All GLMs were set up according to FSL’s recommended guidelines for group analysis and details can be found on the <u>FSL – FMRIB Software</u> <u>Library</u> website. All analyses were performed independently per contrast.

Each model produced a z-statistics maps, which were thresholded using a fixed-effect, voxel-wise threshold of z > 1.64 with a cluster-wise correction of z > 1.64 (p-value < 0.05) to identify significantly active voxels. These maps were overlaid on the PAM50 T2w spinal cord template (De Leener et al., 2017) to visualize activation across the cervical spinal cord (Figure 3). Segmental activation distributions were quantified using SCT’s PAM50 spinal segment masks (PAM50_spinal_levels.nii.gz) specifically from the C5 to C8 spinal cord segments (Figure 1C, Figures 3, 4, and 6). To further characterize the spatial distribution of spinal cord activity, two voxel-based ratios were computed. The first was the Gray/White Matter Ratio (GW Ratio), which quantified the relative distribution of significant voxels across GM and WM by dividing the percentage of active GM voxels by those in WM (Weber et al., 2020). The second was the Left/Right Index (LR Index), which measured lateralization by computing the normalized difference between active voxels in the left and right hemicords, and dividing the difference by the sum (Weber et al., 2020; Weber et al., 2016b).

**Figure 3.**
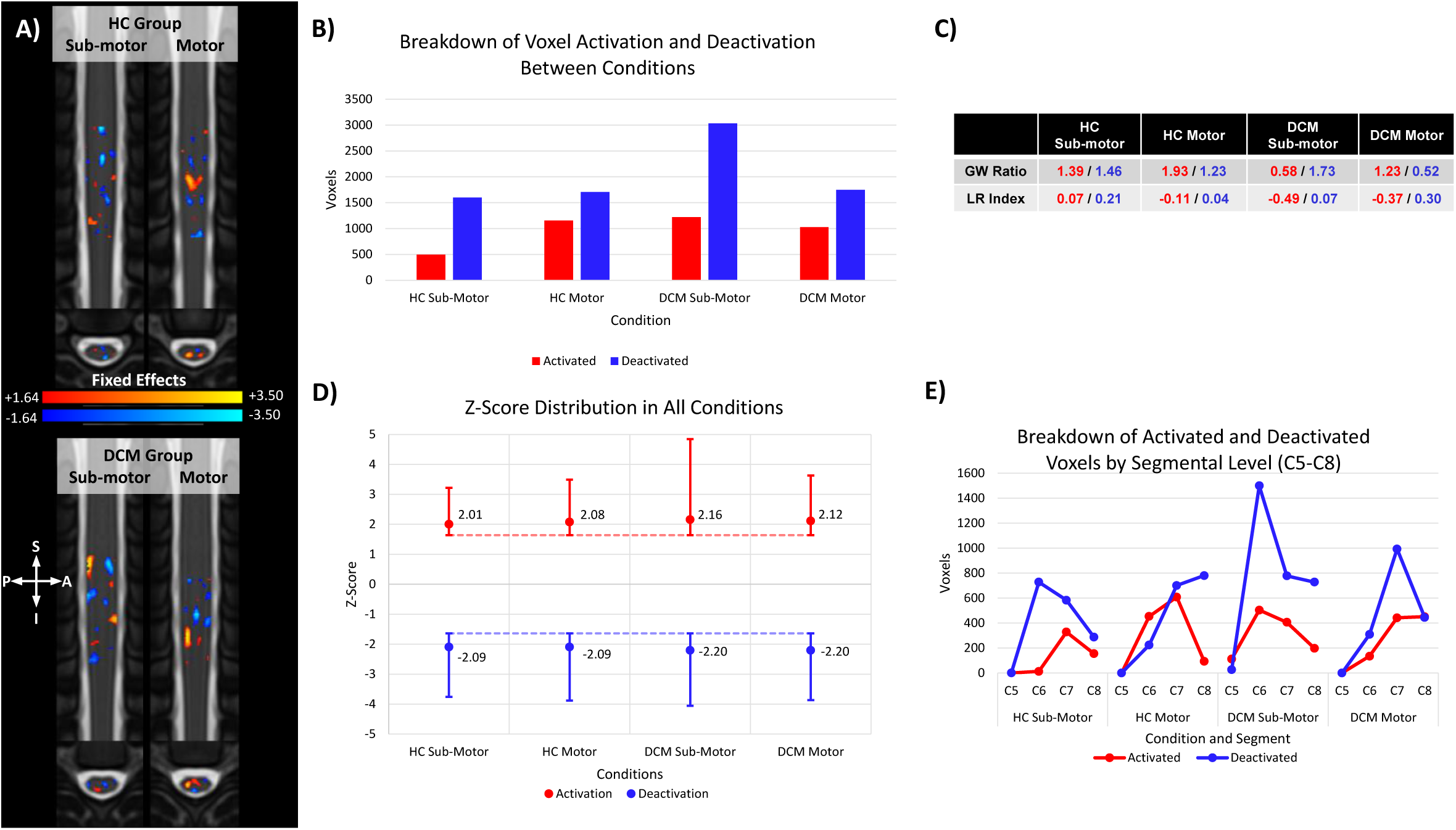
Spinal cord task-based fMRI BOLD response in healthy controls and patients with DCM. (A) Fixed-effects activation maps for sub-motor and motor conditions in healthy controls (HC, top) and degenerative myelopathy patients (DCM, bottom). Activations (red-yellow) and deactivations (blue-light blue) are displayed on axial and sagittal spinal cord T2w (PAM50) templates spanning cervical segments C5–C8. Cluster thresholds are set at ±1.64 to ±3.50. (B) Group-level Voxel count summary showing the number of significantly activated (red) and deactivated (blue) voxels for each condition and group. (C) Summary table showing the gray matter (GM) to white matter (WM) activation ratios (GW Ratio) and left-right laterality indices (LR Index) across conditions. (D) Average Zscore distributions for activated (red) and deactivated (blue) voxels across all conditions, with standard error bars. (E) Segmental-level analysis (C5–C8) of voxelwise activation and deactivation, indicating condition-specific and segmental variations in BOLD signal in DCM and HC.

## 3 Results

### 3.1 Spinal Cord fMRI Responses to Sub-motor and Motor Median Nerve Stimulation

To identify regions with statistically significant BOLD activation (BOLD z-statistic 0) and deactivation (BOLD z-statistic < 0), we performed a one-sample t-test at the group level for HCs and DCM groups. Significant BOLD responses were observed in both HC and DCM during sub-motor and motor threshold stimulations (Figure 3A and Supplemental Figure 2). In the HC group, motor stimulation elicited greater activation across the C5-C8 segments (1155 voxels) compared to sub-motor stimulation (496 voxels), while deactivation remained relatively consistent across both conditions (1598 vs 1705 voxels, Figure 3B). In contrast, DCM patients showed robust changes in deactivation between sub-motor and motor conditions (3032 vs 1747 voxels), while activation remained relatively stable (1220 vs 1028 voxels, Figure 3B).

Spatial distribution in the HC group was predominantly localized to the GM in both sub-motor and motor conditions (GW Ratio: sub-motor = 1.39; motor = 1.93) (Figure 3C). However, the DCM group showed a reverse in GM localization during the sub-motor condition (0.57), but aligned more with HCs during motor stimulation (1.23) (Figure 3C). Furthermore, lateralization patterns (LR index) in HCs were relatively balanced (sub-motor: 0.07; motor: -0.11), whereas the DCM group showed right-sided lateralization (sub-motor: -0.49; motor: -0.37) (Figure 3C). Across the two conditions, the mean z-scores for activation was similar between groups (HCs: sub-motor = 2.01, motor = 2.08; DCM: sub-motor = 2.16, motor = 2.12) (Figure 3D). The z-scores for deactivation were also comparable (HC: sub-motor = -2.09, motor = -2.09; DCM: sub-motor = -2.20, motor = -2.20) across both conditions (Figure 3D).

The segmental level (C5-C8) activation revealed condition-specific differences. In the HC group, motor stimulation elicited caudal increasing activation from C6 (453 voxels) to C7 (608 voxels) (Figure 3E). Sub-motor responses were reduced across all segments showed (C6: 13 voxels, C7: 328 voxels, C8: 155 voxels) compared to their motor counterparts, with exception of C8 (Figure 3E). In DCM patients, activation decreased caudally from C6 (504 voxels), C7 (406 voxels), and C8 (198 voxels) segments during sub-motor stimulation and increased slightly during motor stimulation (C6: 135 voxels, C7: 442 voxels, C8: 451 voxels) (Figure 3E). Notably, the motor activation in the DCM group was lower than in HC group across most spinal segments, particularly at C6 and C7. Deactivation in HCs had a caudal decrease across C6 (728 voxels), C7 (583 voxels), and C8 (287 voxels), but increased across the same segmental levels (C6: 224 voxels, C7: 701 voxels, C8: 780 voxels) during motor stimulation (Figure 3E). Deactivation in DCM was elevated in C6 (1500 voxels), C7 (778 voxels), and C8 (728 voxels) segments during sub-motor. During motor stimulation, deactivation responses decreased, with exception of C7, (C6: 309 voxels, C7: 993 voxels, C8: 445 voxels), but still remained higher when compared to HC, with exception of C8 (Figure 3E). Additionally, during sub-motor stimulation, the DCM group showed both activation (112 voxels) and deactivation (26 voxels) at C5, which was not observed in the HC group (Figures 3E).

### 3.2 Group Level Changes Between Sub-motor and Motor Stimulation

To determine the changes in spinal cord activation across stimulation intensities, we performed a paired, two-sample t-test comparing motor and sub-motor stimulations in each group. Significant activation differences were detected in all comparisons (motor sub-motor and sub-motor > motor; Figure 4). In the HC group, motor stimulation evoked a greater increase in activation compared to sub-motor stimulation, activation with a 4.09% difference in active voxels during motor vs 2.12% during sub-motor (i.e. difference in active voxels divided by number of total voxels in the C5-C8 mask) (Figure 4B).

**Figure 4:**
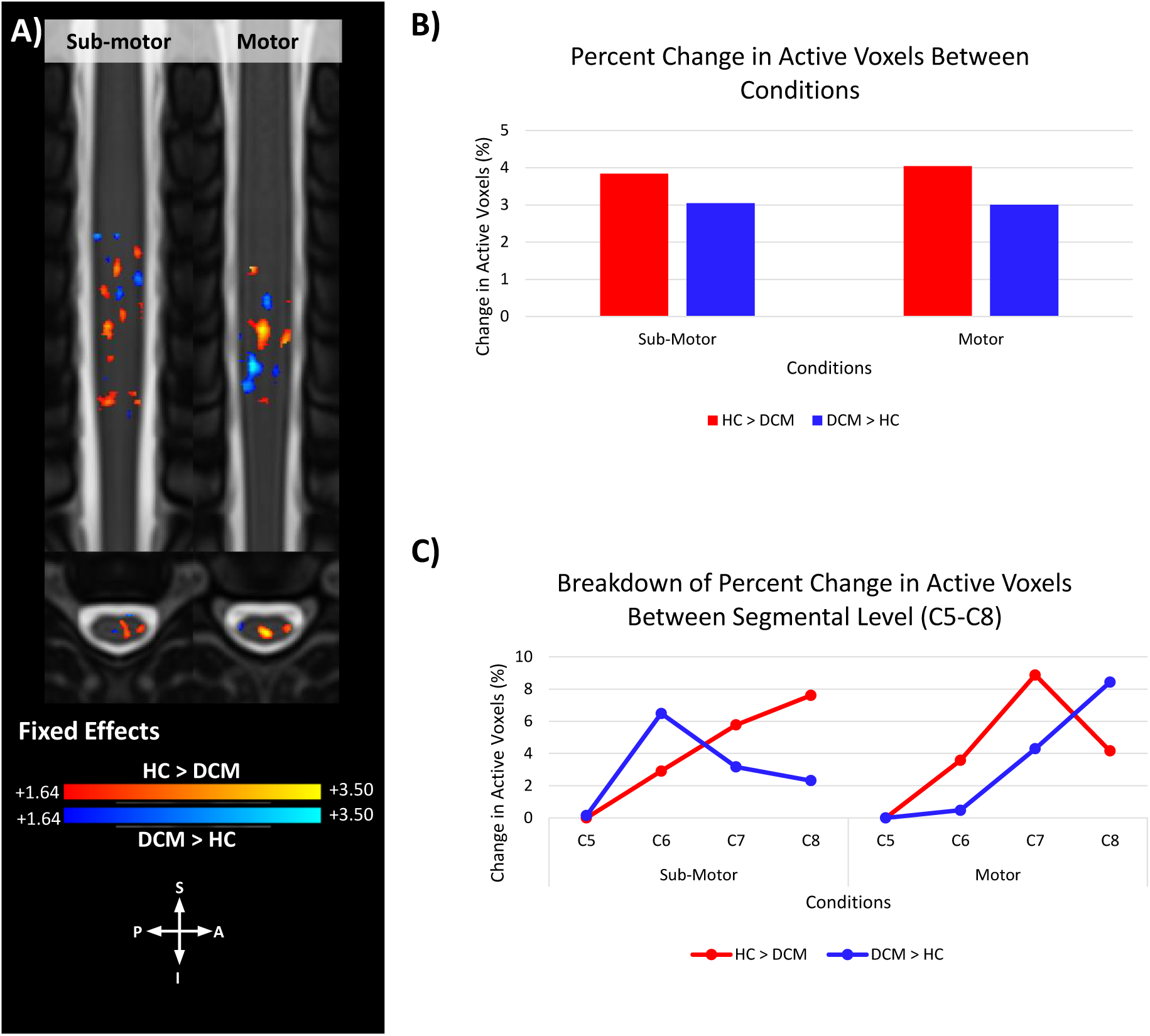
Motor and sub-motor threshold contrasts-driven BOLD responses in HC and DCM. (A) Fixed effects activation maps of threshold response showing where motor > sub-motor (red-yellow) and sub-motor > motor (blue-light blue) responses in healthy controls (HC, left) and degenerative myelopathy patients (DCM, right). Activations maps are overlaid on T2 w axial and sagittal (PAM50) template. Separate comparison maps are shown for HC and DCM groups. (B) Percent change in the number of active voxels between motor and sub-motor conditions within each group (C) Segmental-level (C5–C8) distribution of active voxels in motor > sub-motor (red) or sub-motor > motor (blue) contrasts.

Different patterns of segmental response were observed between HC and DCM due to changes in the stimulation intensities. In the HC group, the comparison of motor sub-motor revealed the largest increase in activation at C6, followed by a decrease caudally (C6: 9.41%; C7: 5.35%; C8: 1.22%). In contrast, the sub-motor > motor comparison showed a caudal increase in activation with more voxels at lower segments (C6: 1.15%; C7: 3.44%; C8: 4.53%), suggesting that the sub-motor threshold preferentially recruited lower spinal segments (Figure 4C). The DCM group exhibited a different response pattern. The sub-motor > motor comparison revealed larger increase in activation overall than the motor > sub-motor comparison. Specifically, sub-motor > motor comparison showed increasing activation from C6 (5.39%) to a peak at C7 (8.89%), then C8 (6.86%). In contrast, the motor > sub-motor response was weaker across all segments (C6: 3.23%; C7: 7.11%; C8: 2.57%) (Figure 4C). This divergent pattern indicates that an increase in stimulation intensity does not enhance spinal cord activation in DCM patients and that responses to motor and sub-motor stimulations may be less distinct than in HC.

### 3.3 Voxel Activation at Subject Level

To further examine individual response and variability within each group, we analyzed subject level changes in the number of active voxels between sub-motor and motor conditions (Figure 5A). More HCs had a modest increase in active voxels from sub-motor to motor stimulation (13/23) in HC group compared to 5/23 in the DCM group (Figure 5A). Furthermore, patients in the DCM group showed a more inconsistent response with majority showing reduced activation during motor stimulation (18/23). On average, the DCM group exhibited a decrease of 212 voxels when transitioning from sub-motor to motor stimulation, while the HC group exhibited a modest positive shift in mean voxel counts (17.83 voxels) (Figure 5B). The DCM group also showed a greater inter-subject variability and a negative shift (Figure 5B). When the percent changes were calculated, HCs again showed a small increase in activation (7.06%), whereas DCM patients exhibited an overall decrease (-2.23%) (Figure 5C). However, these differences at the subject level were not statistically significant.

**Figure 5:**
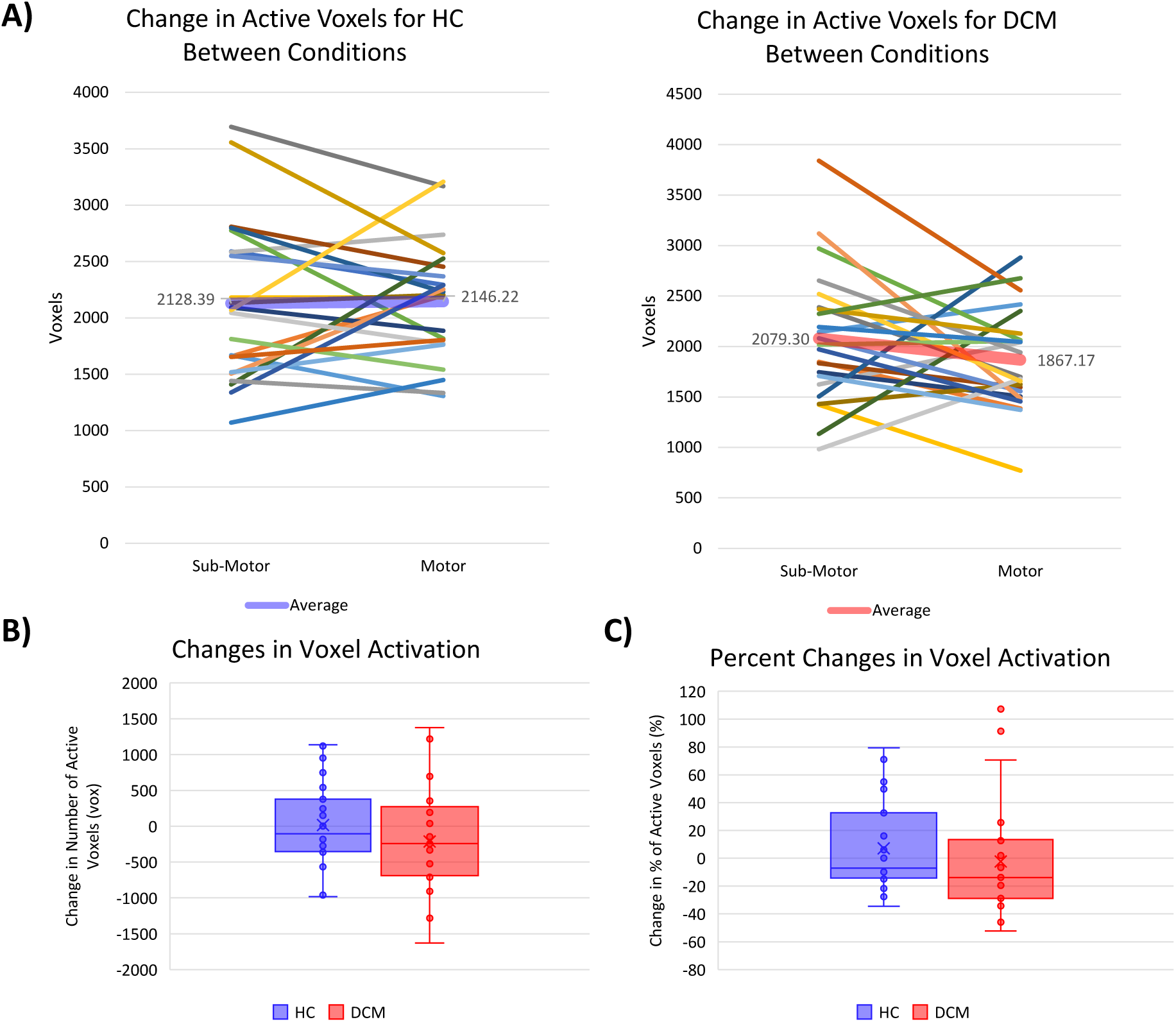
Subject-level changes in spinal cord activation between motor and sub-motor conditions in HC and DCM. (A) Line plots showing individual changes in active voxel counts between sub-motor and motor conditions for each subject, in healthy control (HC, left) and degenerative myelopathy (DCM, right) groups. Mean voxel change between the conditions are overlaid in color (HC: blue and DCM: red). (B) Boxplot of the absolute differences in voxel activation per subject between conditions (motor minus sub-motor) for each group (C) Boxplot of percent difference in active voxels between conditions per subject. Box plot, *HC: blue, DCM: red*.

### 3.4 Group Level BOLD Changes Between HC and DCM

While the earlier analyses characterized responses within each group, here we directly compared HC and DCM activation patterns using unpaired, two-sample t-test between (HC > DCM; DCM > HC) for both sub-motor and motor stimulation conditions. This analysis allowed us to quantify and spatially map group-level differences (Figure 6A). Overall, HCs showed greater difference in the amounts of active voxels during sub-motor and motor stimulations over DCM between groups (3.84% and 4.04% respectively, with the difference in active voxels being divided by the number of total voxels in the C5-C8 mask) (Figure 6B).

**Figure 6:**
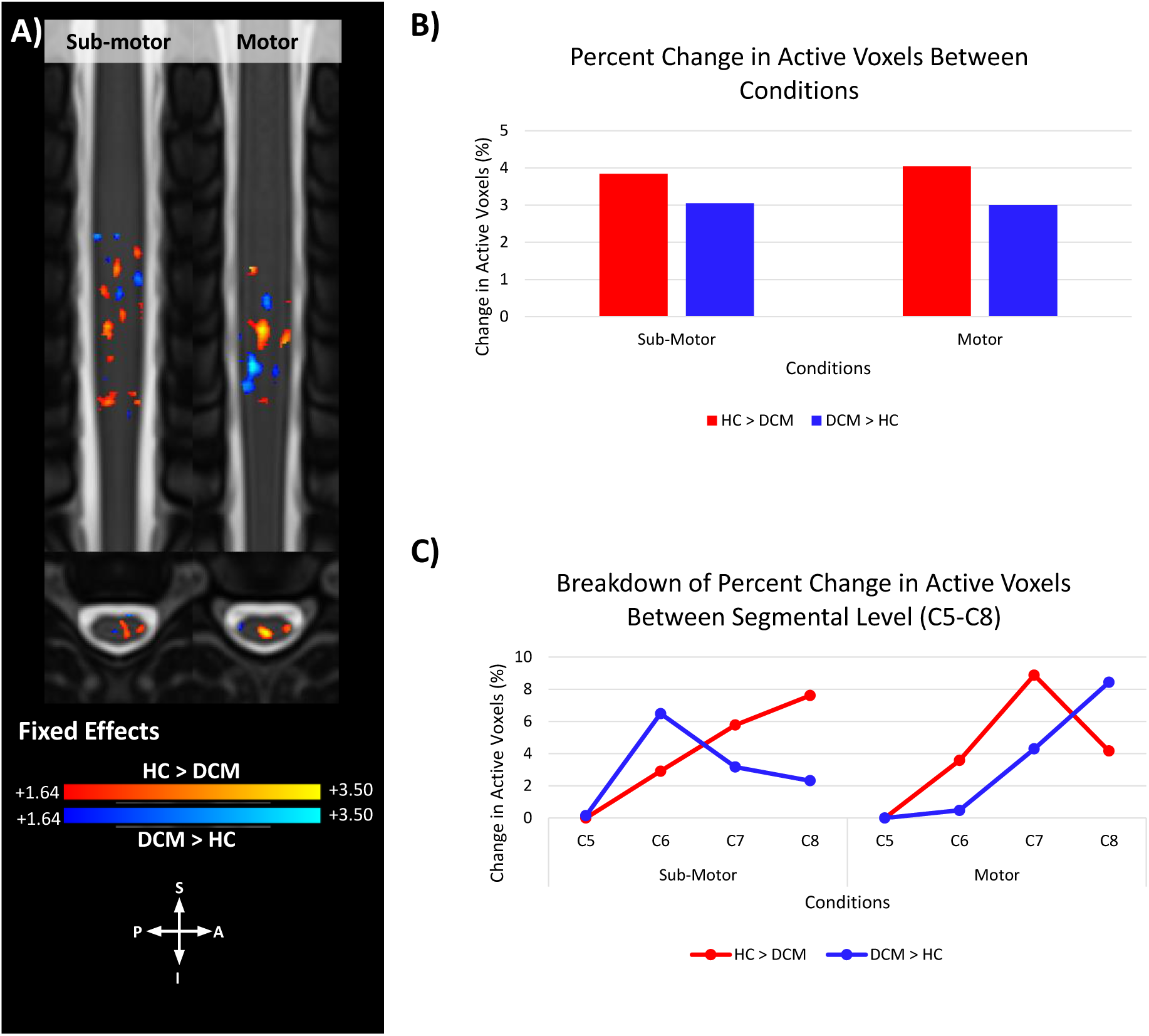
Group level differences comparing HC and DCM across the sub-motor and motor conditions. (A) Fixed-effects contrast maps showing voxel-wise differences between HC and DCM groups during sub-motor and motor tasks. Red-yellow indicates HC > DCM, while blue-light blue indicates DCM > HC. BOLD activation maps overlaid on axial and sagittal T2w PAM50 templates. (B) Percent change in the number of significantly active voxels by group and condition derived from unpaired two sample t test. Red bars is activation in HC > DCM, while blue bars reflect DCM > HC. (C) Segmental-level breakdown of differential activation (C5–C8), showing voxel-wise segmental group difference.

Segmental level comparisons (Figure 1C) provided additional insight into these group differences. In HCs, motor stimulation exhibited the greatest increase in activation than DCM at C7 (8.88%) (Figure 6C). In contrast, the DCM-HC pattern showed a progressive caudal increase (C6: 0.47%; C7: 4.3%; C8: 8.43%), peaking at C8, and suggesting a shift in segmental response between the two groups. Furthermore, the HC-DCM difference peaked at C8 (7.61%) during sub-motor stimulation, whereas the DCM-HC pattern shifted rostrally peaking at C6 (6.48%) (Figure 6C). These suggest that DCM patients exhibited enhanced sub-motor responses at slightly more rostral segment compared to motor stimulation, which adds to the evidence indicating alterations in the localization of spinal cord activation in DCM.

## 4 Discussion

Degenerative cervical myelopathy can cause progressive damage to both the white matter (WM) and gray matter (GM) pathways of the spinal cord (Cloney et al., 2018; Haynes et al., 2024; Hopkins et al., 2018; Smith et al., 2020). However, while WM and structural injury have been well studied (Bedard et al., 2025; Cloney et al., 2018; Filimonova et al., 2024; Haynes et al., 2024; Muhammad et al., 2024; Paliwal et al., 2020), less is known about the functional consequences of GM injury, despite its role in sensory and motor output. Defining these changes is clinically important because a greater injury burden predicts poorer surgical outcomes (Kopjar et al., 2018; Tetreault et al., 2015). In this study, we apply a task-based fMRI paradigm using peripheral median nerve stimulation, a method that helps to avoid reliance on voluntary motor performance to provide a more consistent measure of spinal cord functional activity.

In healthy controls, our group level analysis of spinal cord activation shows an expected increase between sub-motor and motor conditions. This enhanced activation response to an increase in stimulation is expected, as a greater stimulation intensity will recruit motor neurons that have a higher activation threshold than sensory neurons (Gaines et al., 2018). Furthermore, the strongest recruitment was seen at C6 and C7, which aligns with the spinal nerve root origins of the median nerve and electrophysiological studies showing that stimulation results in a higher amplitude/current in the vertebral levels aligning with the C7/C8 segments (Akaza et al., 2021). However, DCM patients showed a paradoxical decrease in BOLD activated voxels when the stimulation intensity and abnormal recruitment at C6 and C8 during sub-motor stimulation. This expansion of activation suggests impairments in localized neuronal recruitment and the presence of compensatory reorganization in DCM. Sub-motor stimulation responses were also exaggerated relative to motor, pointing to abnormal excitability or impaired habituation of compressed pathways. This excessive sub-motor response was specifically pronounced in the C6 segment, increasing by compared to its HC counterpart, which aligns with the vertebral levels commonly associated with DCM compression (i.e. C5 and C6) (Muhammad et al., 2024; Vallotton et al., 2021). Finally, activation lateralization, determined by the LR index, appeared to be primarily bilateral in HCs. Ipsilateral responses in the spinal cord are expected because the neurological signals involved in the spinal cord reflex arc enter and exit on the ipsilateral side. Contralateral activation (i.e. the side opposite of the stimulation) is also consistent with intraneuronal connectivity cross at the spinal cord level and has been previously reported in other task-based fMRI paradigms (Agosta et al., 2009; Bouwman et al., 2008; Brooks et al., 2012; Maieron et al., 2007; Ng et al., 2008). DCM responses, however, were predominantly ipsilateral to the stimulation, implying that, in addition to the segmental shift in activation, DCM patients also experience a decrease in intraneuronal/left-right connectivity that is preventing contralateral activation.

In addition to regions of positive BOLD signal change (i.e. activation), regions of negative BOLD signal change (i.e. deactivation) were noted in all conditions. In the brain, deactivation has long been suspected to be the result of inhibition (Allison et al., 2000; Nakata et al., 2019; Schafer et al., 2012). This indicates that changes in deactivation patterns could also be important because they reflect additional alterations in neuronal processes. Deactivation has also been reported to extend to the spinal cord, where it inhibited neuronal activity in the left ventral and dorsal horns during a motor task (Oliva et al., 2025). These findings in the spinal cord align with the known presence of inhibitory interneurons, which are known to help regulate the transmission of sensory signals (Bardoni et al., 2013; Goulding et al., 2014), especially pain (Hughes & Todd, 2020), and motor reflex coordination (Goulding et al., 2014). In our data, deactivation was observed to be consistent in HCs across stimulation intensities and showed a bilateral distribution (i.e. GW Ratio). In contrast, DCM patients exhibited a marked spike at C6 during sub-motor stimulation, far greater than its HC counterpart. This abnormal inhibitory signal coincided with increased activation and higher z-scores in sub-motor stimulation, suggesting that heightened excitation and inhibition co-occur at the same segment. Notably, this phenomenon was localized to the C6 segment (C5 vertebral level), the site most affected in DCM (at C6 during sub-motor stimulation, the segment most compressed in DCM (Muhammad et al., 2024; Vallotton et al., 2021). This exaggerated inhibitory response could be a reflection of a maladaptive processing of the excitatory drive at low threshold and underscores the role of inhibitory interneurons in shaping abnormal activity pattern (Bardoni et al., 2013; Goulding et al., 2014; Hughes & Todd, 2020). Beyond indicating areas of deactivation, negative BOLD signals have been increasingly recognized as biologically meaningful. Altered deactivation patterns during spinal cord stimulation have been associated with pain processing in surgical spine patients (De Groote et al., 2018) and linked to inhibitory interneuron activity regulating sensory transmission and motor reflexes (Oliva et al., 2025). In combination with our own data, these studies help to support the idea that heightened inhibition in DCM reflects disrupted sensorimotor integration.

The resulting change in segmental and lateral cord recruitment among DCM participants displayed by direct stimulation is likely reflective of the structural damage occurring during cord compression. Previous biomechanical DCM studies have shown that diffused compression on the anterior part of the cord causes the most stress (Levy et al., 2021) and results in the most detrimental symptomatic outcomes (Nishida et al., 2012). This stress can then cause disruption to the capillaries of the blood-spinal cord barrier, ultimately causing ischemia (i.e. lack of blood flow) to the spinal cord (Badhiwala et al., 2020; Blume et al., 2020; Tu et al., 2021) that can impair neural function through several mechanisms. The first is that the damage has caused neuronal loss and impeded activation, which has been documented in the WM and GM (Badhiwala et al., 2020; Blume et al., 2020; Cloney et al., 2018; Haynes et al., 2024; Haynes et al., 2025; Hopkins et al., 2018; Smith et al., 2020; Tu et al., 2021) and indicates the presences of ventral horn atrophy and motor neuron degeneration. These structural changes align with clinical reports of motor performances deficits in DCM patients (Muhammad et al., 2023; Muhammad et al., 2024). The second possible explanation is that surviving neurons remain structurally intact but functionally impaired. Compressive DCM damage that results in axonal degeneration and demyelination (Akter et al., 2020; Dohle et al., 2023) will reduce conduction and responsiveness and therefore change functional responses in the spinal cord. In response, axons can reroute or recruit alternative segments, a similar reaction to spinal cord injury (Sofroniew, 2018), which can result in reorganization of neuronal pathways and an abnormal spread of activation to the C5 and C8 segments. Finally, the ischemia may blunt BOLD signal sensitivity by reducing the amount of oxygenated blood being delivered to active regions (Glover, 2011).

Together, these processes can provide a better insight into the pathophysiological mechanisms for the functional alterations we observed in DCM patients during task-based fMRI.

The ability to detect functional changes in DCM patients with fMRI highlights the advantages of a median nerve stimulation paradigm. Backes et al., was the first to demonstrate that median nerve stimulation could evoke cervical spinal cord activation and was comparable to voluntary motor tasks such as fist clenching (Backes et al., 2001). Subsequent work confirmed that median nerve stimulation can produce robust BOLD responses in the cervical spinal cord (Xie et al., 2007) validating the paradigm as a reproducible method to probe sensorimotor pathways. The HC group also displayed activation profiles aligning with electrophysiological depictions of median nerve stimulation in the cervical spine (Akaza et al., 2021), supporting the validity of the paradigm’s consistency. Furthermore, this approach presents the ability to avoid dependence on patient motor performance, which is a critical limitation of task-based DCM studies due to motor impairments (Muhammad et al., 2023), and revealed distinct functional alterations in DCM patients. This underscores the paradigm’s potential as a method to assess spinal cord function in disease state.

Because of this, applying a direct median nerve stimulation fMRI paradigm in DCM patients could provide one of the first functional biomarkers to complement established structural imaging. In the past, structural MRI has helped to define cord compression, atrophy, and other degenerative changes (Bedard et al., 2025; Cloney et al., 2018; Filimonova et al., 2024; Haynes et al., 2024; Khan et al., 2023; Muhammad et al., 2024; Smith et al., 2020). However, it cannot capture functional activity. By utilizing techniques that rely on detecting changes in blood oxygenation (i.e. BOLD signal) within spinal cord segments affected by DCM, we demonstrate an approach that offers insight into segmental and lateral recruitment patterns and inhibitory alterations that are not accessible with structural imaging. The ability to indirectly map the functional changes within the regions of damage highlights the potential median nerve stimulation has potential to serve as a clinically relevant, functional DCM biomarker.

### 4.1 Limitations

There are several limitations to this study. First, spinal cord compression can distort cord anatomy and vertebral alignment, which may affect template registration and the normalization to the PAM50 template that is derived from healthy controls (De Leener et al., 2018). Secondly, spinal cord fMRI has known challenges with reproducibility, particularly in response to stimulation where signal levels are typically lower than during voluntary motor tasks (Backes et al., 2001; Wilmink et al., 2003). This may contribute to variability across individuals and limit the generalizability of group-level comparison. Third, stimulation thresholds were determined based on observed finger twitch, which is inherently subjective and may introduce variability in application. While pilot work with EMG has confirmed reflex responses for this paradigm, future studies should incorporate objective electrophysiological measures to improve consistency. We also observed that all DCM patients required higher motor thresholds to elicit comparable responses (Haynes et al., 2025), suggesting that there are disease-related differences in excitability that must be accounted for in future refinements of this technique.

### 4.2 Future Directions

The data presented here demonstrates functional changes in the spinal cords of DCM patients during sub-motor and motor stimulation. However, little is known about reorganization of resting-state patterns in DCM. While dorsal and ventral resting-state networks have been defined in healthy controls (Barry et al., 2014; Kong et al., 2014), evidence in DCM is limited to a single study associating BOLD fluctuations with JOA scores (Liu et al., 2016). Examining resting-state patterns and comparing them to task-based stimulation responses could reveal whether the extended activation we observed at C5 and C8 represents maladaptive dysfunction or adaptive reorganization in response to stimulation. Furthermore, because DCM is a progressive disease that can be classified by severity (Tetreault et al., 2017), future studies should investigate whether functional changes differ between mild and severe categories (i.e., moderate and severe). Such comparison could improve understanding of DCM progression and may inform decisions on surgical intervention and predict outcomes, thereby positioning spinal fMRI as a prognostic tool of DCM. Lastly, an important next step will be to establish how these functional changes relate to clinical motor and sensory deficits in DCM. Linking functional activation and deactivation patterns to symptom severity could clarify the contribution that abnormal excitability and reorganization have to disability.

Further work should also examine how functional changes couple with established structural alterations, providing insight into how anatomy and physiology interact to shape disease progression.

## 5 Conclusion

In summary, direct median nerve stimulation combined with fMRI revealed functional alterations in DCM patients, which included abnormal segmental recruitment, exaggerated sub-motor responses, reduced motor activation, altered lateralization, and increased inhibitory processing at compressed levels. These changes likely reflect compensatory mechanisms that allow partial motor function to be preserved despite tract damage. By capturing the functional disruptions not visible with structural imaging, this paradigm highlights a potential avenue for developing functional spinal cord biomarkers of DCM.

## 6 Data and Code Availability

As this work is part of an ongoing longitudinal study, the complete dataset is not publicly available at this time. However, de-identified imaging data can be made available upon reasonable request to the senior author.

## 7 Author Contributions

**Grace Haynes:** Methodology, Investigation, Writing – Original Draft, Writing – Review and Editing, Visualization; **Fauziyya Muhammad:** Methodology, Software, Formal analysis, Investigation, Writing – Original Draft, Writing – Review and Editing; **Kenneth A. Weber 2nd:** Conceptualization, Methodology, Writing – Review and Editing; **Ali F. Khan:** Investigation, Writing – Review and Editing; **Sanaa Hameed:** Investigation; **Hakeem J. Shakir:** Resources; **Michael Rohan:** Resources; **Lei Ding:** Methodology, Writing – Review and Editing, Supervision; **Zachary A. Smith:** Conceptualization, Methodology, Resources, Writing – Original Draft, Writing – Review and Editing, Project Administration, Supervision, Funding Acquisition

## Acknowledgements

The authors would like to acknowledge the contributions of the MRI technicians and staff at the Laureate Institute of Brain Research (LIBR) in Tulsa, OK, who helped with data acquisition.

---

**Supplemental Figure 1:**
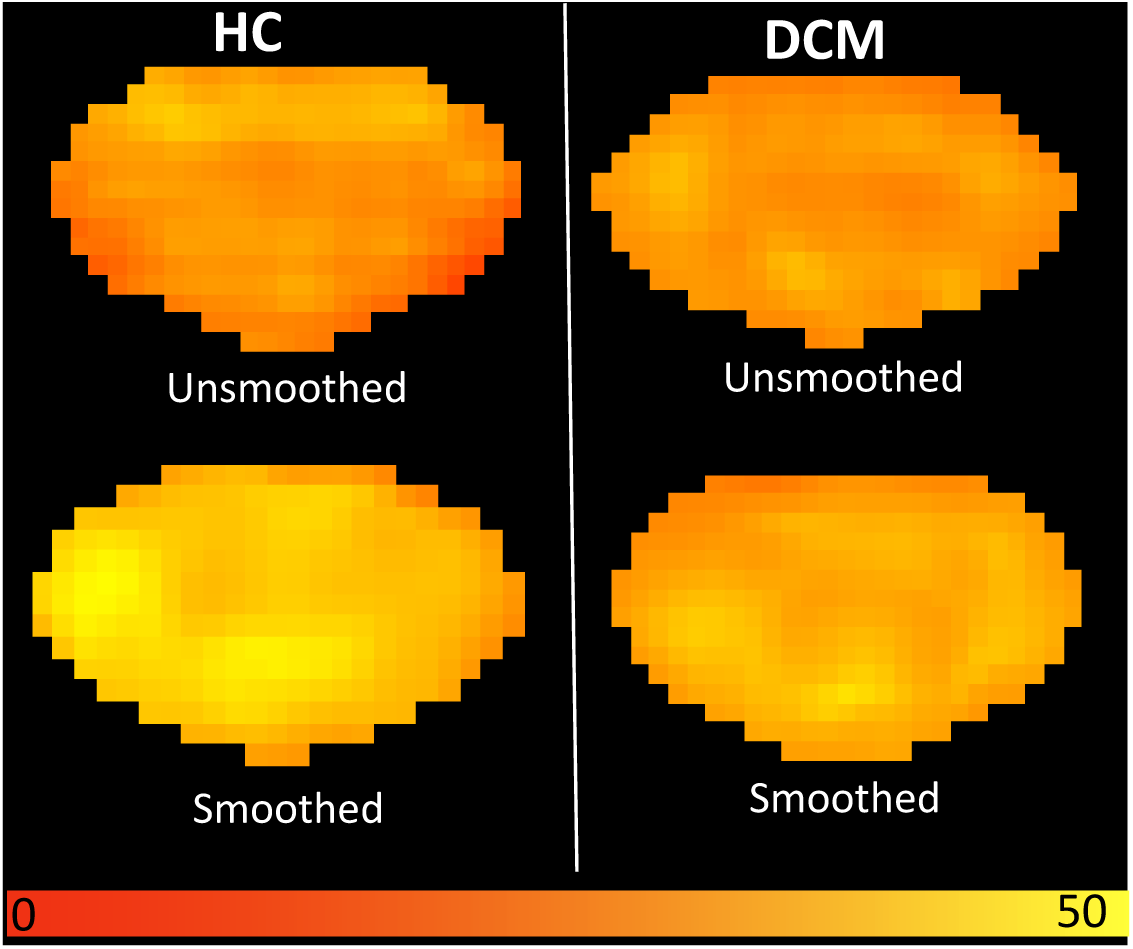
Average spinal cord temporal signal-to-noise ratio (TSNR) maps from a representative healthy control (HC, left) and degenerative myelopathy (DCM, right) subject at C5 vertebral level.

**Supplemental Figure 2:**
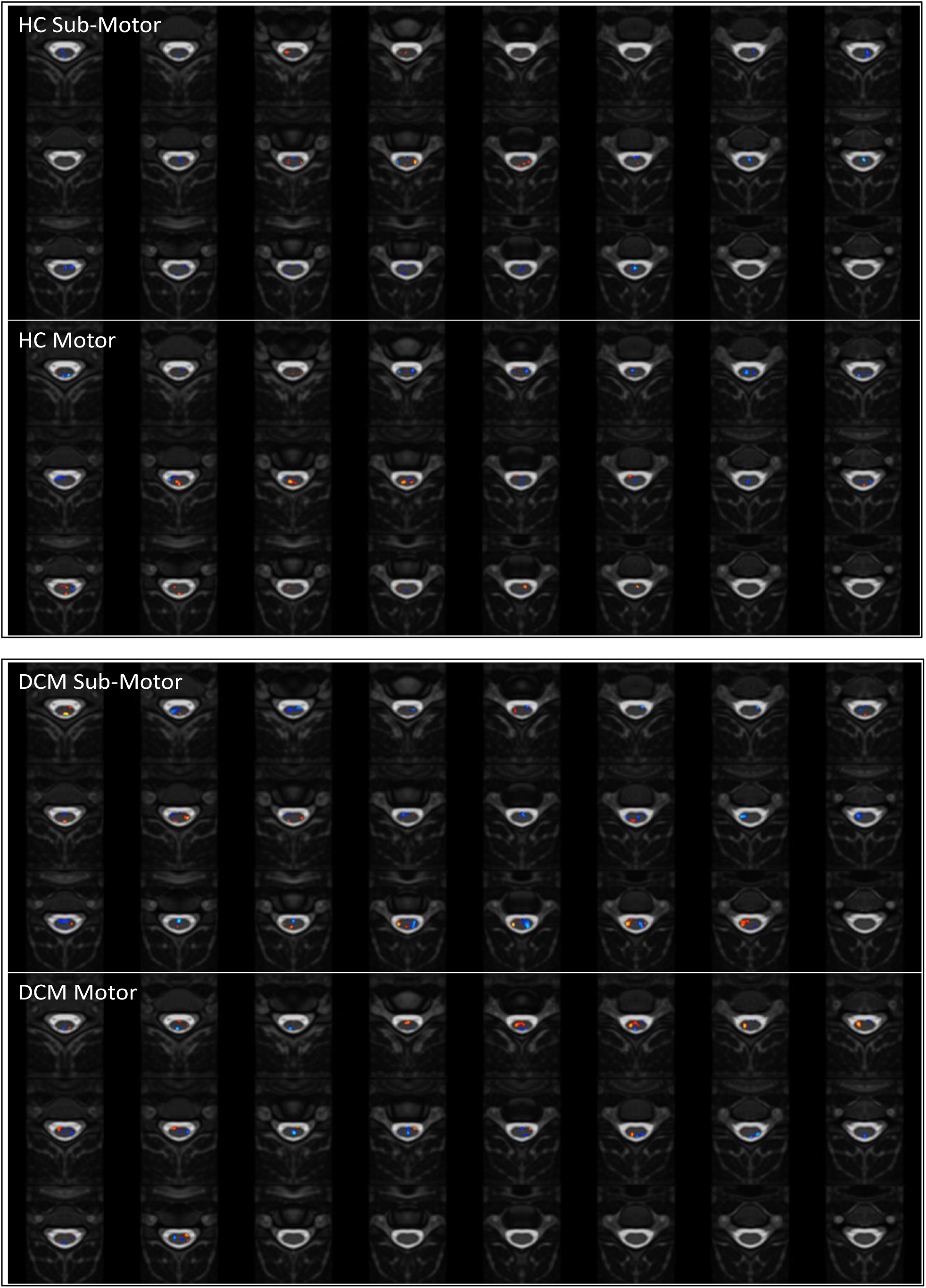
Axial slices over the C4-C7 vertebra displaying activation (red) and deactivation (blue) in healthy controls (HC, top) and degenerative myelopathy (DCM, bottom) groups.

